# Human placental syncytiotrophoblasts restrict *Toxoplasma gondii* vertical transmission at two distinct stages and induce CCL22 in response to infection

**DOI:** 10.1101/170944

**Authors:** Stephanie E. Ander, Elizabeth N. Rudzki, Nitin Arora, Yoel Sadovsky, Carolyn B. Coyne, Jon P. Boyle

**Author notes:** Address correspondence: Carolyn Coyne, PhD, 9116 Rangos Research Center, Pittsburgh, PA 15260, Phone (412) 695-, Jon Boyle, PhD, 4249 Fifth Avenue, Pittsburgh, PA. 15260.

## Abstract

*Toxoplasma gondii* is a major source of congenital disease worldwide, but the cellular and molecular factors associated with its vertical transmission are largely unknown. In humans, the placenta forms the key interface between the maternal and fetal compartments and forms the primary barrier that restricts the hematogenous spread of microorganisms. Here, we utilized primary human trophoblast (PHT) cells isolated from full-term placentas and human mid-gestation chorionic villous explants to determine the mechanisms by which human trophoblasts restrict and respond to *T. gondii* infection. We show that placental syncytiotrophoblasts, multinucleated cells that are in direct contact with maternal blood, restrict *T. gondii* infection at distinct stages of the parasite lytic cycle—at the time of attachment and also during intracellular replication. Utilizing comparative RNAseq transcriptional profiling, we also show that human placental trophoblasts at both mid- and late-stages of gestation induce the chemokine CCL22 in response to *T. gondii* infection, which relies on the secretion of parasite effector(s). Collectively, our findings provide new insights into the mechanisms by which the human placenta restricts the vertical transmission of *T. gondii* at early and late stages of human pregnancy, and demonstrate the existence of at least two interferon-independent pathways that restrict *T. gondii* access to the fetal compartment.

**Significance statement:** *Toxoplasma gondii* is a major source of congenital disease worldwide and must breach the placental barrier to be transmitted from maternal blood to the developing fetus. The events associated with the vertical transmission of T. gondii are largely unknown. Here, we show that primary human syncytiotrophoblasts, the fetal-derived cells that comprise the primary placental barrier, restrict *T. gondii* infection at two distinct stages of the parasite life cycle and respond to infection through the induction of the chemokine CCL22. Collectively, our findings provide important insights into the mechanisms by which human syncytiotrophoblasts restrict *T. gondii* infection at early and late stages of human pregnancy and identify the placental-enriched signaling pathways induced in response to infection.

## Introduction

*Toxoplasma gondii* is a major source of congenital disease, with ~200,000 global cases of congenital toxoplasmosis reported each year (1). In the majority of instances (~80%), *in utero* infections by *T. gondii* result in a range of severe birth defects, including ocular disease and developmental delays, and can also result in fetal death (2). However, despite the clear impact of *T. gondii* infections on fetal health, the mechanisms by which the parasite is transmitted from the maternal bloodstream into the fetal compartment are largely unknown.

In eutherian organisms, the placenta serves as the sole source of gas, nutrient, and waste exchange from the fetal compartment and acts as a key barrier to restrict fetal infections. At the forefront of these defenses is the syncytiotrophoblast (SYN), a multinucleated cell layer that comprises the outermost layer of the human placenta and which is in direct contact with maternalperfusion of the intervillous space. Subjacent to the SYN layer are cytotrophoblasts (CYTs), mononucleated and proliferative cells that fuse to replenish the SYN layer throughout pregnancy. Together, these trophoblast layers form a key barrier to the passage of pathogens that may infect the fetus by the hematogenous route.

In general, the pathways that exist in the human placenta to limit the vertical transmission of microbes are poorly defined. Our previous studies in primary human trophoblast (PHT) cells, which were focused on viral pathogens, have identified at least two potent antiviral pathways that restrict viral replication in trophoblasts (3, 4). However, these pathways do not appear to be relevant during infection with non non-viral pathogens, including *T. gondii* (5). While studies in placental explants suggest that the SYN layer is not permissive to *T. gondii* infection (6), the mechanistic basis for SYN resistance is incompletely understood, as is whether the SYN layer mounts any innate defense in response to parasite exposure. Moreover, while placental explant models are useful in their recapitulation of placental structure, they are limited in their capacity to dissect trophoblast cell type-specific pathways that might exist to limit *T. gondii* infection.

In this study, we interrogated the trophoblast cell-type specificity of *T. gondii* infection utilizing PHT cells isolated from full-term placentas and identified two cellular mechanisms that mediate SYN-specific resistance to *T. gondii* infection. In addition to discovering that SYNs restrict *T. gondii* attachment and replication, we also identified cell signaling pathways uniquely induced by parasite infection in PHT cells, which included the release of the regulatory T cell (Treg) chemoattractant CCL22. We show that the majority of transcriptional changes in PHT cells is specific to *T. gondii* and does not occur in response to infection with the closely related parasite *Neospora caninum*. Moreover, we show that CCL22 induction is dependent on host cell invasion and the secretion of *T. gondii* effectors into the host cell. To expand our findings to earlier in human pregnancy, when the fetus is usually more susceptible to congenital *T. gondii* infections, we also utilized a mid-gestation chorionic villous explant model and show that second trimester SYNs also resist *T. gondii* attachment and induce CCL22 in response to infection whereas the fetal-derived amnion and chorion are permissive to infection and do not induce CCL22. Taken together, we have identified previously unknown intrinsic features in primary human placental cells from both the second and third trimesters of pregnancy that limit *T. gondii* infectivity at the level of invasion and replication, and provide details on both host- and parasite-specific transcriptional responses of placental cells to infection.

## RESULTS

### Syncytiotrophoblasts isolated from term placentas resist *T. gondii* infection

We found that PHT cells isolated from full term placentas exhibited reduced susceptibility to *T. gondii* infection when compared to primary human foreskin fibroblast (HFF) cells (**Supplemental Figure 1A, 1B**). These data are consistent with our previous work demonstrating that PHT cells exhibit reduced susceptibility to infection by the three major types of *T. gondii* in North America and Europe compared to non-placental cells (7). Importantly, human trophoblast cell lines (including BeWo, HTR8, and JEG-3 cells) were unable to recapitulate this restrictive phenotype and were permissive to parasite infection (**Supplemental Figure 1C**). In addition, this phenotype was specific to PHT cultures as primary placental fibroblasts were as permissive to infection as HFF cells (**Supplemental Figure 1D**).

PHT cells isolated from full-term placentas spontaneously fuse to form SYNs during their culture period (~72hrs), with some retaining a mononuclear CYT phenotype. Therefore, to determine whether the lack of PHT cell infection occurred in a cell-type specific manner, we infected PHT cells with YFP-tagged *T. gondii* (RH strain) and quantified parasite growth specifically in CYTs versus SYNs. These studies revealed dramatic differences in the susceptibility of SYNs and CYTs to *T. gondii* infection—whereas CYTs were permissive to infection, SYNs were highly resistant (**Figure 1A, left**). Furthermore, we observed that parasites within SYNs replicated to a lesser degree, as indicated by a highly significant reduction in total cell area occupied by parasites (**Figure 1A**). Since *T. gondii* replicates within a parasitophorous vacuole (PV) generated at the time of invasion by each individual parasite, the number of parasites within a PV can serve as an indicator of parasite growth and replication. In contrast to the PVs in CYTs, those in SYNs most often contained 1-2 parasites (**Figure 1A, right**). Importantly, fusion of BeWo cells with forskolin, which induces syncytin-mediated fusion (7), was not sufficient to confer resistance to *T. gondii* infection (**Supplemental Figure 1E**), supporting that this phenomenon is specific to primary cells.

**Figure 1.**
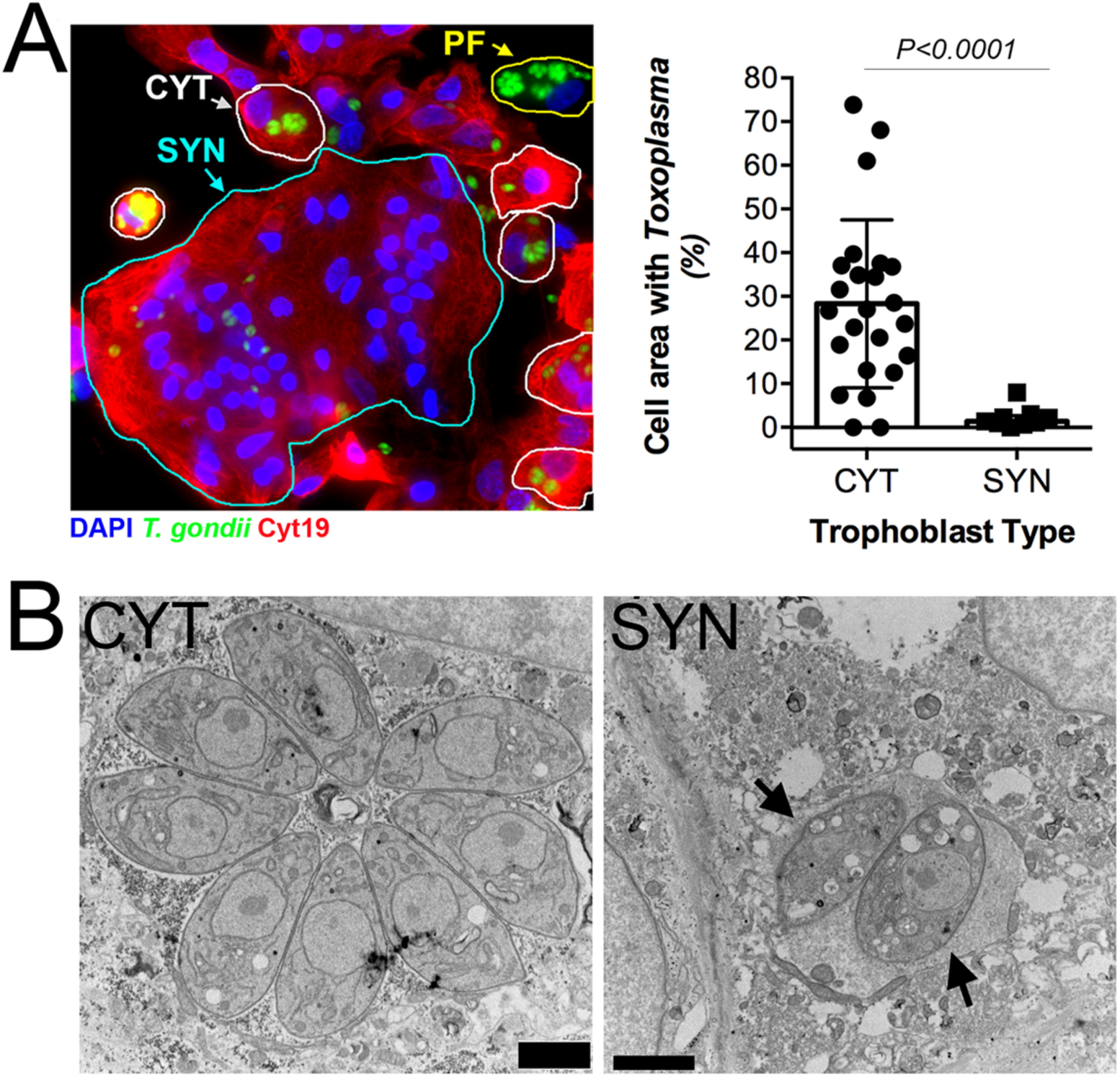
Placental syncytiotrophoblasts resist *T. gondii* infection. **(A)** Left, immunofluorescence microscopy of PHT cells inoculated with *T. gondii* RH strain (green) for ~24h. Cytokeratin-19 is shown in red; DAPI in blue. SYN (outlined in cyan), CYT (outlined in yellow), and placental fibroblast (PF, outlined in white). PF cells are distinguished by the lack of cytokeratin-19 (red). Right, percentage of cell area occupied by *T. gondii*, as compared between CYT and SYN. N_CYT_=24, N_SYN_=9. 2-tailed T-test *P*<0.0001 with Welch correction for unequal variances. **(B),** Transmission electron microscopy of PHT cells infected with *T. gondii* (RH) for ~23hpi. Mononucleated CYT at left and SYN, identified by its more than two nuclei at right. Scale bar is 2 µm.

Transmission electron microscopy (TEM) revealed that whereas parasite growth and PV morphology were normal in mononucleated cells within the preparation (which are likely CYTs but could also be rare contaminating placental fibroblasts), SYN-internalized parasites were found within PVs containing host cell cytoplasmic contents indicative of a loss of vacuole integrity (**Figure 1B**). Moreover the parasites within these PVs contained more vacuoles of minimal electron density and poorly defined organelles (**Figure 1B**). This phenotype is reminiscent of drug-induced death that we observed previously after treatment with a benzodioxole-containing compound (8), indicating that SYNs have potent Toxoplasmacidal activity. Taken together, these data implicate SYNs as an innately resistant cellular barrier to *T. gondii* infection and suggest that unlike other cultured cells, these cells potently resist *T. gondii* infection.

### SYN-mediated resistance to *T. gondii* infection occurs at two stages of the parasite lytic cycle

Our data suggest that SYNs restrict *T. gondii* infection at a stage of intracellular parasite growth. The primary mechanisms for cell-autonomous immunity to *T. gondii* are driven by the effector cytokine interferon γ (IFNγ). However, we found that uninfected and *T. gondii*-infected PHT cells had low levels of IFNγ transcript (**Figure 2A**) and that culture supernatants were devoid of secreted IFNγ protein (**Figure 2A, 2B**). Importantly while the expression of GBP1 and GBP2 as well as other innate immunity-related factors (e.g., NOS1,2 and IDO) have comparatively higher transcript levels in PHT cells (**Figure 2A**), none of these well-characterized IFNγ-driven host effector proteins were uniquely expressed in PHTs (**Figure 2A**). These findings suggest that the innate resistance of SYNs is not dependent on basal expression of IFNγ or its stimulation by infection. To explore other related mechanisms, we performed immunofluorescence microscopy for markers of autophagy- and lysosomal-mediated degradation pathways given the high level of basal autophagy previously observed in PHT cells (9). We found that there was no association between SQSTM/p62 or the lysosomal associated component LAMP2 and internalized parasites at either early or later stages of infection (**Figure 2C** and **Supplemental Figure 2)**. These findings are consistent with our TEM-based microscopic studies, in which we also did not observe the association between double membraned autophagosomes or lysosomes with internalized parasites (**Figure 1B**).

**Figure 2.**
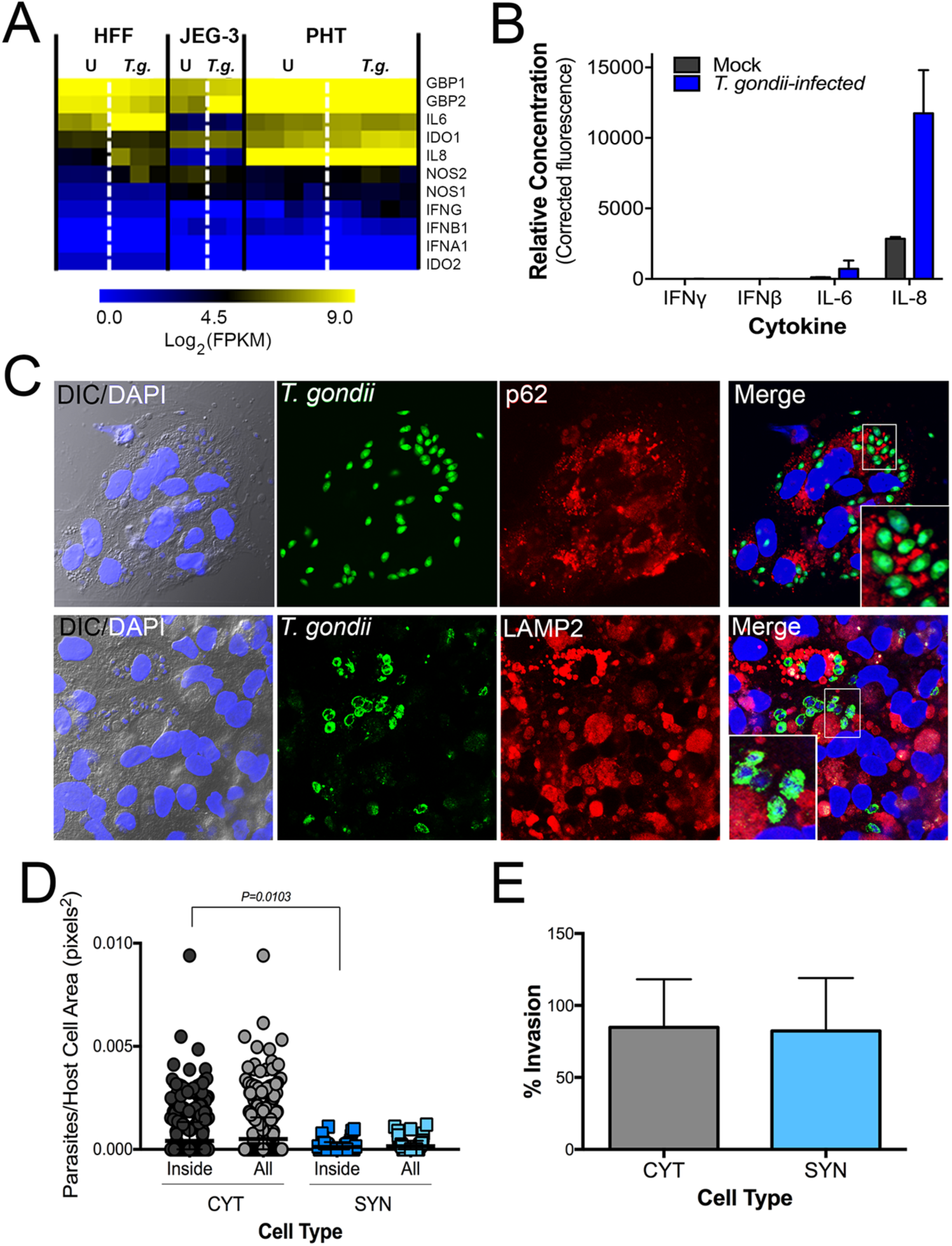
SYN-mediated restriction of *T. gondii* infection is not the result of autophagy, lysosomal degradation, or inability to invade. **(A)** Heat-map of innate immune effector gene expression as determined by RNAseq of uninfected and *T. gondii-*infected HFF, JEG-3, and PHT cells. **(B)** PHT cytokine secretion as detected by luminex assay of media from mock and *T. gondii-*infected PHT cells. **(C)** Immunofluorescence microscopy of PHT cells infected with *T. gondii* (RH) (green) at 8hpi. *(Top)* Infection with YFP-RH at MOI=10; p62 staining is shown in red. *(Bottom)* Infection with RH (anti-GRA2, green) at MOI=2; lysosome-associated membrane protein 2 (LAMP2) is shown in red; DAPI in blue **(D, E)** Quantification of inside/outside staining of *T. gondii* (YFP-RH) infected PHT cells at 2hpi, MOI=1. Using DIC/DAPI images, cells were classified as mononucleated (CYT) or multinucleated (SYN). (D) To compare the attachment efficiency between cell types, the number of internalized parasites per 2D cell area was calculated for 813 cells (N_CYT_=729, N_SYN_=84) and the resulting distributions were compared using the Kolmogrov-Smirnoff test. The comparison used was: CYTinside vs SYNinside (*P*=0.010). (E) Comparison of percent invaded parasites of all parasites-associated cells by cell type ((N_CYT_=176, N_SYN_=34).

In addition to the intracellular control of parasite replication, it is possible that SYNs are protected from infection by defects in parasite attachment and/or invasion. To quantify parasite attachment and invasion, we performed a two-step immunofluorescence-based invasion assay to distinguish extracellular from intracellular parasites (as in Kafsack et al. (10) and others). PHT cells were exposed to *T. gondii* (RH-YFP) for 2 hrs, at which point monolayers were washed to remove unbound parasites, cells were fixed, and attached parasites were identified using an antibody against surface antigen-1 (SAG1) in the absence of cell permeabilization, followed by detection using a secondary antibody conjugated to Alexa Fluor 594. Samples were then permeabilized and incubated again with anti-SAG1 antibody, which was detected with a secondary antibody conjugated to Alexa Fluor-633. Using this approach, extracellular parasites exhibit fluorescence in all channels (YFP, 594, and 633) whereas intracellular parasites exhibited fluorescence in only two (YFP and 633). Using differential contrast imaging (DIC) and automated image analysis, we quantified the extent of attached and internalized parasites in CYTs versus SYNs, which were easily distinguishable using DIC based upon the number, size, and clustering of their nuclei. Using this approach, we found that there were significantly fewer parasites overall (i.e., uninvaded and invaded) that were associated with SYNs compared to CYTs (normalized for cell area; p=0.010; **Figure 2D**). However, the percentage of invasion events (of all total parasite associations) was nearly identical between SYNs and CYTs, demonstrating that while there is a significant defect in parasite attachment to and/or association with, SYNs, there is no obvious impediment to invasion (**Figure 2E**). While we do not know the stage of attachment at which *T. gondii* tachyzoites are arrested when associating with SYNs compared to CYTs, these data point to a defect in *T. gondii* attachment as a primary mediator of SYN resistance to infection in addition to their ability to potently resist parasite replication.

### PHT cells have a unique response to *T. gondii* infection characterized by the induction of immunity-related transcription factors and chemokines

Given the dramatic differences in infectivity and growth of *T. gondii* in PHT cells, we infected PHT cells or the choriocarcinoma JEG-3 cell line with *T. gondii* and compared their transcriptional responses to infection using RNAseq. Following infection for 24 h, we identified 401 transcripts of significantly different abundance (P<0.01; Fold-change>4) in infected PHT cells, and 106 transcripts of different abundance in infected JEG-3 cells (**Figure 3A and 3B**). To identify which transcripts were uniquely induced in PHT cells compared to JEG-3 cells and another primary cell line, we compared these data to a recently published RNAseq dataset from *T. gondii*-infected primary HFF cells(11). While we identified 858 host cell transcripts that were of different abundance in *T. gondii*-infected HFFs, there was a significant lack of overlap between infection altered transcripts in HFF and PHT cells (**Figure 3B**). Cluster analysis of all genes induced in *T. gondii*-infected PHT cells revealed multiple categories of genes specifically induced in these cells and not in either HFFs or JEG-3 cells. While some genes were induced uniquely in PHT cells and were expressed poorly in other cell types, (**Figure 3A**, cluster “a”), others were of high abundance only after infection in PHT cells, but constitutively expressed in other cell lines/types (**Figure 3A**, cluster “b”). Focusing on genes uniquely induced in PHTs compared to other cell types (“Cluster 1”, **Figure 3C**), multiple immunity-related transcription factors (e.g., IRF4, EGR4), chemokines (CCL22, CCL17, CCL20, CCL1) and the chemokine receptor CCR7 were all significantly induced by *T. gondii* infection. In particular, we found that CCL22, a chemokine known to be expressed constitutively during pregnancy(12, 13) and that has also been found to increase during miscarriage(12) was induced by >400 fold in infected PHT cells based on RNAseq (**Figure 3C**), which was confirmed in independent PHT preparations using RT-qPCR (>1000-fold; **Figure 3D,** left) and at the protein level by ELISA on infected PHT supernatants (from ~25 pg/mL in mock controls to >500 pg/mL in infected cells; **Figure 3D,** right). Heat-killed *T. gondii* failed to induce CCL22 release from PHT cells, indicating that production of this chemokine requires live parasites and suggesting that the CCL22 response requires parasite invasion.

**Figure 3:**
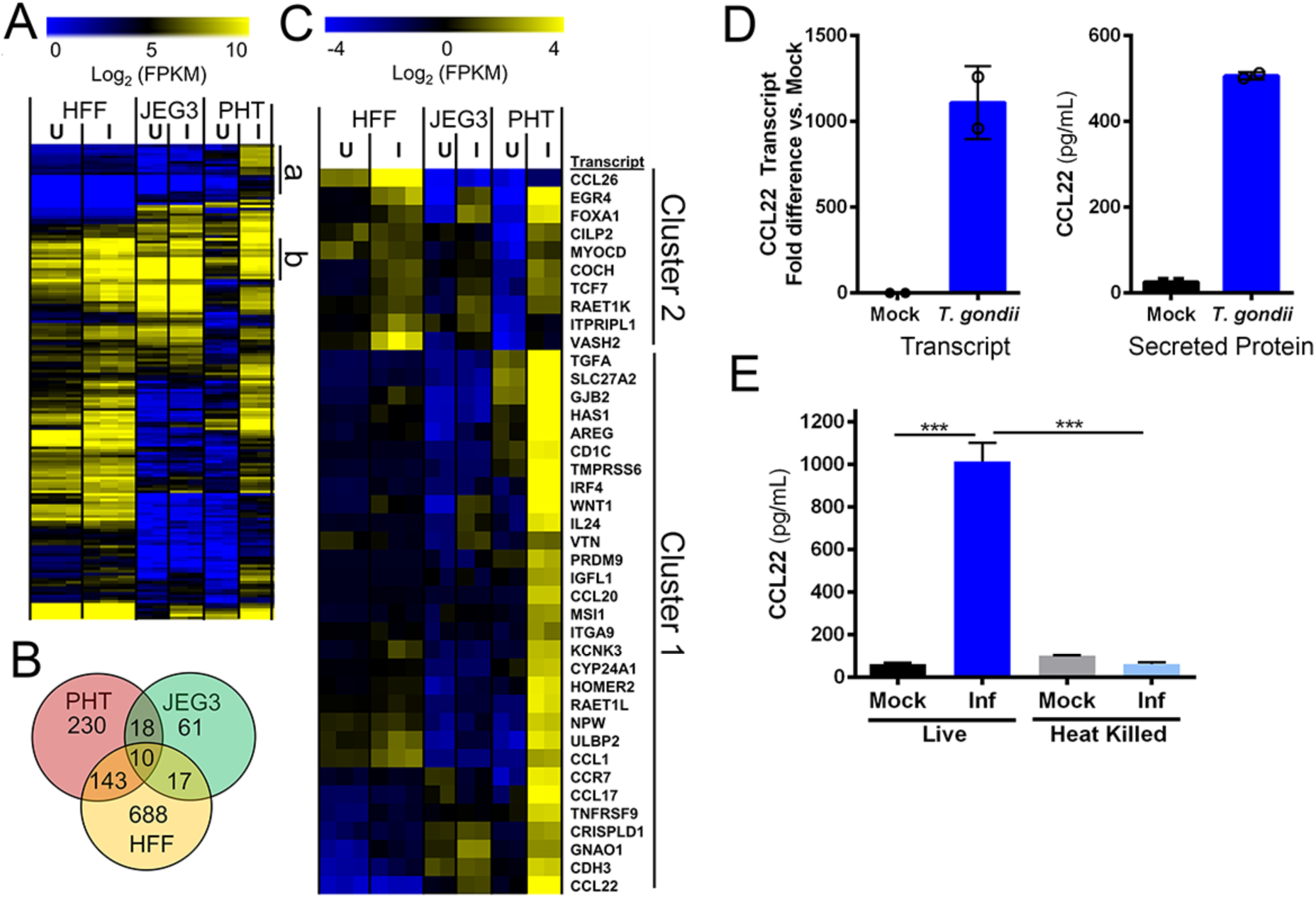
PHT cells infected with *Toxoplasma gondii* have a unique transcriptional response to infection. A) Heat map of all genes with significantly higher transcript abundance in *T. gondii* infected PHT cells compared to mock (P<0.01; Fold-change >4). B) Genes with significantly higher (P<0.01; fold-difference >=4) in PHT cells, HFFs and JEG3s. C) Hierarchically clustered heat map of 40 genes induced in infected PHT cells. Cluster 1 contains genes that are induced in other cell types, while cluster 2 consists primarily of transcripts induced by *T. gondii* infection only in PHT cells (27/30). Transcription factors and chemokines and their receptors are indicated with asterisks. D) CCL22 transcript (left) and secreted protein (right) levels in *T. gondii*-infected PHT cells. N=2 for each assay. E) Induction of CCL22 secretion in PHT cells requires live parasites. Parasites were incubated at 23 \degrees C or 65 \degrees C for 1 h prior to being used to infect PHT cells. ***:*P*<0.001 following 1 way ANOVA and multiple comparisons post-hoc tests.

### Infection of PHT cells with *Neospora caninum* does not induce inflammatory signaling

Host transcriptional responses to infection with *T. gondii* have been shown in a variety of cell types to be specific for *T. gondii* and are not associated with infection by one of its apicomplexan relatives, *Neospora caninum*(14),(15, 16). Unlike *T. gondii*, *N. caninum* is not a human pathogen, but causes significant mortality in cattle and dogs and is associated with congenital disease in these animals(17, 18). To determine the specificity of the host response to *T. gondii* infection in PHT cells, we infected cells with *T. gondii* (RH-YFP) or *N. caninum* (NC-1-dsRED(19)) and compared the cellular responses to infection using RNAseq. We found that *N. caninum* failed to significantly induce any of the chemokine/chemokine receptor genes that were induced by infection with *T. gondii* of PHT cells (**Figure 4A**; asterisks indicate focus chemokine genes). The remarkable lack of differential transcript abundance in *N. caninum*-infected PHT cells compared to matched *T. gondii* infected PHTs was further illustrated by MA plot (**Figure 4B**). In PHT cells, we identified 206 genes that were significantly induced by *T. gondii* infection (P<0.05; fold-induction>2), and only 10 genes that were significantly induced after *N. caninum* infection (**Figure 4C**). Consistent with this, infection with *N. caninum* had no effect on CCL22 levels and co-infections with *N. caninum* and *T. gondii* showed that there was also no synergistic or additive effect (**Figure 4D**). Importantly, despite the significant differences in gene induction, we found that PHT cells were similarly susceptible to *N. caninum* infection, with CYTs being readily invaded and supportive of parasite growth, while SYNs exhibited reduced invasion and growth restriction very similar to that observed for *T. gondii* (**Figure 4E**).

**Figure 4:**
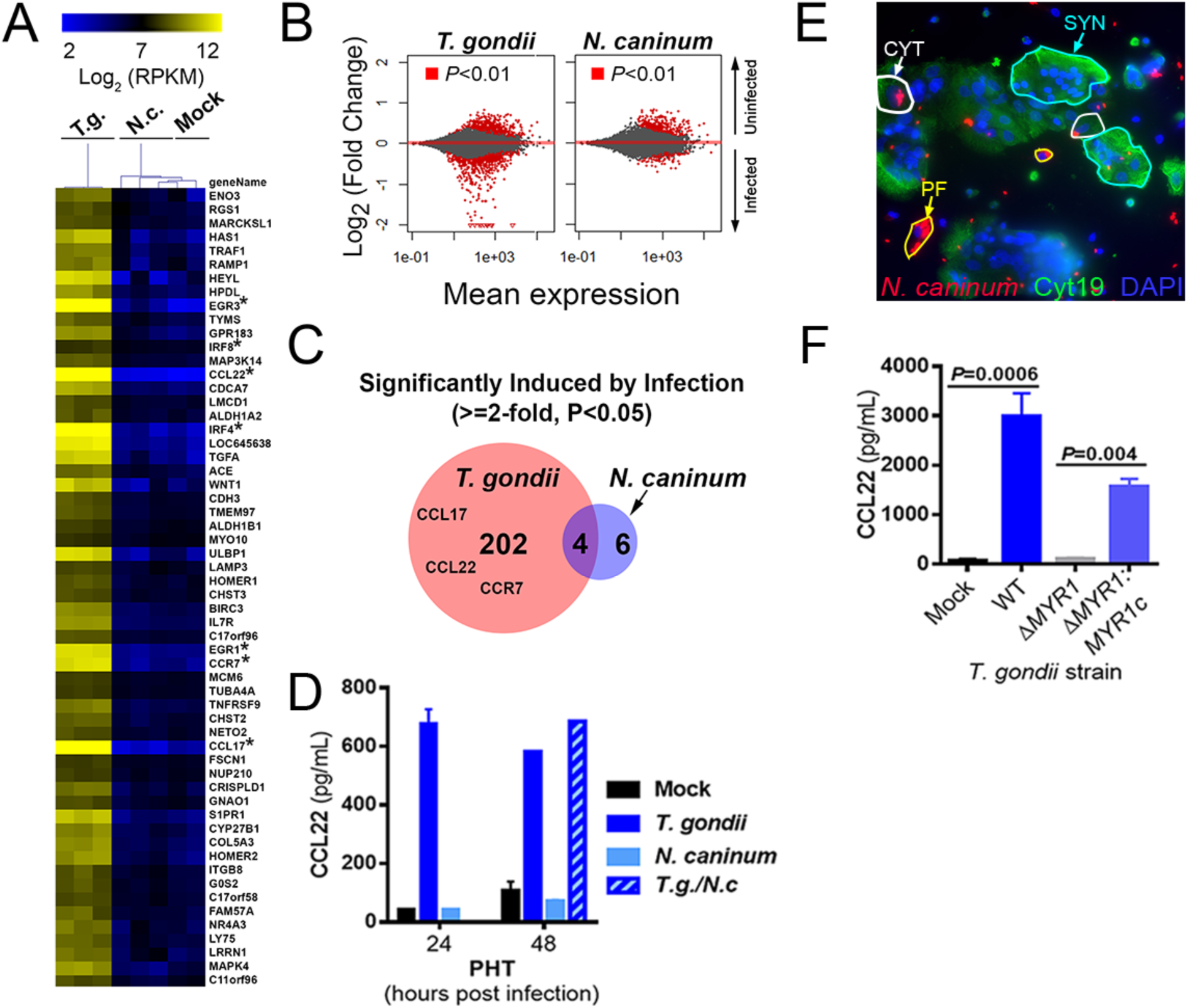
Infection-modulated gene expression is specific to *T. gondii* and requires parasite effectors. A) Heatmap of 59 genes induced by at least 1.8-fold in PHT cells (*P*<0.01) after infection with *T. gondii* RH strain (MOI=3). Raw count data were converted to normalized FPM using DESeq2. Data are also shown for *N. caninum*-infected (N=3) and Mock-infected (N=2) PHTs, and mean centered data were clustered by sample and gene using the Euclidean Distance (implemented in MeViewer; TM4 microarray suite; See Methods). B) MA plots of PHTs comparing gene expression profiles in Mock and parasite-infected cells for all 23,735 queried genes. Genes of higher abundance in uninfected cells are indicated by positive changes and those of higher abundance in infected cells are indicated by negative changes. C) In *T. gondii* infected PHTs, 206 genes were found to be of higher abundance compared to Mock-infected cells, while only 10 such genes were found in *N. caninum*-infected PHTs, consistent with the MA plots in B above. D) ELISA showing induction of CCL22 secretion in PHTs infected with *T. gondii*, but not *N. caninum*. Host cells were infected with an MOI of 2 (for each parasite species) and supernatants were harvested at the indicated time points. N=2-3 wells for all treatments except for 48 h *T. gondii* and *T. gondii*/*N. caninum* (N=1). E) PHTs were infected with NC1:dsRED *N. caninum* (MOI=3) for 24 h, and stained with cytokeratin 19 antibodies and DAPI. Similar to *T. gondii*, *N. caninum* grew efficiently in PFs (yellow outlines), CYTs (white outlines) and poorly or not at all in SYNTs (blue outlines). F) CCL22 induction in PHT cells is requires MYR1. PHTs were infected with either wild type RH:YFP *T. gondii*, RHΔMYR1, or RHΔMYR1 complemented with an HA tagged copy of MYR1 (RHΔMYR1c). N=3 for each *T. gondii* strain.

### CCL22 induction in PHT cells requires the *T. gondii* dense granule protein MYR1

Given that CCL22 induction in PHT cells required live parasites and was not induced by *N. caninum*, we reasoned that CCL22 induction was likely the result of a *T. gondii*-specific parasite effector that would be secreted after host cell invasion. *T. gondii* MYR1 is a recently identified dense granule protein that is required for the export/secretion of multiple dense granule effectors (including GRA24, GRA25 and GRA16;(15, 20, 21)) and was discovered based on its role in mediating *T. gondii*-specific activation of the transcription factor c-Myc(14). When we infected PHT cells with *T. gondii* lacking MYR1 (RH:Δ*MYR1*) and complemented control parasites (RH: Δ*MYR1*:*MYR1c*), we found that CCL22 production by PHT cells was entirely dependent upon MYR1 (**Figure 4F**), which provides strong evidence that *T. gondii* CCL22 induction in PHT cells is driven by (a) MYR1-dependent secreted effector(s).

### Second trimester human placental villi resist *T. gondii* infection and induce CCL22 in response to infection

Because PHT cells are isolated from term placentas, we next determined whether SYNs from earlier in human pregnancy also resist *T. gondii* infection and induce CCL22. To do this, we utilized second trimester chorionic villous explants, which retain the morphology of human placental villi, including a layer of cytokeratin-19 positive SYNs covering the villi surfaces (**Figure 5A**). Consistent with our findings in PHT cells, and the work of others utilizing first trimester explants (6), we found that second trimester SYNs were resistant to *T. gondii* infection, even when infected with very high numbers of parasites (10^7^) (**Figure 5B, left panel**). This resistance appears to be primarily at the level of parasite attachment as we detected very few internalized parasites in placental villi and most parasites detected appeared to be extracellular (**Figure 5B, left panel, white arrows**). Importantly, unlike placental villi, we found that fetal membrane (amnion and chorion) and maternal decidua supported T. gondii replication (**Figure 5B, middle and right panels**), highlighting the specific resistance of placental villi. In addition, consistent with the work of others utilizing first trimester explants (6), we found that CYTs subjacent to the SYN were permissive to *T. gondii* only when the SYN layer was breached (**Figure 5C**)._ These data show that the SYN layer also forms a barrier to *T. gondii* vertical transmission in mid-gestation.

**Figure 5.**
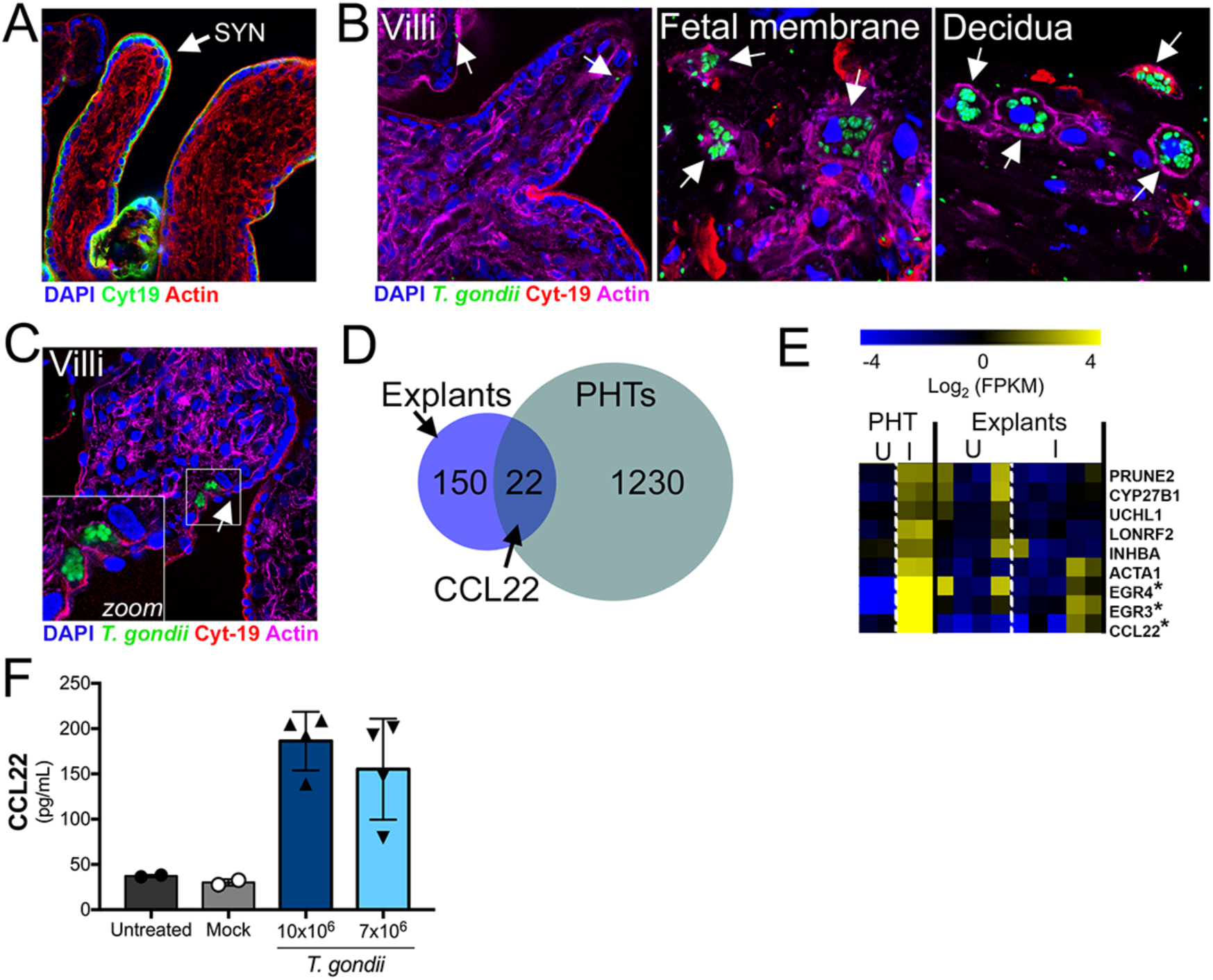
Chorionic villous explants from the second trimester are resistant to *T. gondii* infection and induce similar cytokine profiles in response to *T. gondii* as full-term PHT cells. **(A)** Immunofluorescence microscopy of uninfected second trimester placental villous explant with SYN localization indicated. Cytokeratin-19 in green; actin in red; DAPI in blue. **(B)** Immunofluorescence microscopy of mid-gestation villi (left panel), fetal membrane (middle panel), or decidua (right panel) inoculated with 10^7^ *T. gondii* (RH strain) for 24h. *T. gondii* in green; cytokeratin-19 in red; actin in magenta; DAPI in blue. White arrows denote single parasites in isolated villi and PVs in fetal membrane and decidua. **(C),** Immunofluorescence microscopy of mid-gestation villi inoculated with 10^7^ *T. gondii* (RH strain) for 24h. *T. gondii* in green; cytokeratin-19 in red; actin in magenta; DAPI in blue. Zoomed image from white box denotes PVs in CYTs located beneath a tear in the SYN (white arrow). **(D,E)** RNA-Seq analysis of *T. gondii*-infected villous explants. Venn diagram (D) and heat map (E) showing a subset of genes that are similarly induced in one of the *T. gondii*-infected villous explants (explant 5) and PHTs. Genes were selected if they were at least 2-fold induced compared to mock and significant using a DESeq2-determined adjusted *P*-value<0.05. Explant data in heat map are from 3 genetically distinct placenta preparations, while PHT data shown are the same as in Fig. 3. **(F)** ELISA showing *T. gondii* (RH strain)-mediated enhancement of CCL22 secretion from villous explants at 24hpi.

Next, we profiled the transcriptional changes induced by *T. gondii* infection of second trimester chorionic villi using RNAseq to determine whether they responded to parasite infection similarly to PHT cells from late gestation. We found that 172 transcripts were differentially expressed in response to *T. gondii* infection of villous explants, with 22 of these transcripts also being differentially expressed in response to infection of PHT cells, which included EGR3, EGR4 and CCL22 (**Figure 5D, 5E**). We confirmed that CCL22 was induced at the protein level by ELISA in supernatants from *T. gondii*-infected second trimester villi (**Figure 5F**). These data suggest that CCL22 is specifically induced by the human placenta in response to *T. gondii* infection at both early and late stages of pregnancy.

## DISCUSSION

*T. gondii* infections present a worldwide threat to pregnant women and there is an urgent need to develop novel treatment regimens to block congenital transmission of *T. gondii* and other pathogens that pose risks to the developing fetus. Our data presented here point to a direct role for SYN-intrinsic pathways in the protection of the fetus from *T. gondii* infection, making the SYN a highly unique cell type given that a diverse array of cells studied to date are susceptible to *T. gondii* infection and replication. Our studies in both human primary term and second trimester SYNs suggest that these cells evade *T. gondii* infection at two critical stages of the parasite lytic cycle—at the point of parasite association with the host cell and during intracellular growth. Moreover, by comparing cell-type specific transcriptional profiles from *T. gondii* infected primary trophoblasts and placental tissue with those of other cell types, we identified unique sets of genes induced by *T. gondii* infection in the human placenta, including the induction of CCL22 that requires the presence of parasite-encoded MYR1. Collectively, our data provide significant advances in our understanding of how the human placenta controls, and responds to, *T. gondii* infection.

We found that the first point of SYN-mediated restriction of *T. gondii* infection occurred at the level of parasite association and/or attachment, which we observed both in SYNs isolated from full-term placentas and from mid-gestation chorionic villi. These findings are consistent with the previous work of others suggesting that attachment might be reduced in first trimester SYNs (6), although this was not directly tested. Our attachment and invasion data from PHT cells provide direct evidence that SYNs naturally restrict parasite attachment but are susceptible to invasion once parasite attachment occurs, which appears to be a rare event in SYNs. It remains unclear at what point of the attachment process that is altered when *T. gondii* associates with SYNs, but a likely stage is during the early phase of gliding motility prior to the second phase of attachment mediated by secretion of microneme and rhoptry organelles (22). One possibility is that when parasites encounter SYNs they glide less efficiently to CYTs or placental fibroblasts, which ultimately results in significantly reduced “full” attachment (mediated by microneme and rhoptry secretion). Differences in membrane biochemistry in SYNs versus CYTs and fibroblasts could underlie these important differences in early parasite association. However, once secondary attachment occurs invasion may then proceed normally, when parasites encounter a second level of resistance.

In mid-gestation chorionic villi, the poor association/attachment phenotype was even more profound than that observed in PHT cells, with little to no parasite association with the villi observed. These findings suggest that in addition to biochemical surface differences between SYNs and CYTs, the morphology of the SYN layer itself may directly impact parasite association and attachment. This could be influenced by a number of morphologic differences in this model, including the positive membrane curvature associated with the significant branching of the placental villous trees, which might impact lipid and/or protein composition. In addition, the apical surfaces of SYNs associated with placental explants may be more differentiated than PHT cells, which might impact parasite attachment through the presence of a highly dense brush border in the explant model. Consistent with this, we previously observed very little *T. gondii* attachment in a bead-based three-dimensional cell line model of human SYNs, which also induces significant membrane curvature and allows for the formation of a well-differentiated brush border (7).

For parasites that attach to the SYN layer, our data suggest a second level of resistance to infection that occurs post-invasion. Importantly, in contrast to other cells types, our data show that this resistance is not mediated by IFNγ, which is not basally expressed in PHT cells or induced by *T. gondii* infection. Furthermore, we did not find any evidence for autophagy- or lysosomal-mediated degradation pathways in the intracellular restriction of parasite replication. To date, all known “cell-autonomous” mechanisms of parasite killing in human cells rely on previous stimulation with IFNγ. For example, IFNγ can induce a variety of downstream effector mechanisms depending on the cell type, including tryptophan starvation in HFFs (23, 24), decoration of the vacuole with guanylate binding proteins (25), ubiquitin or other markers for autophagy including LC3B (26, 27). Ultimately, these pathways would lead to lysosomal fusion with the parasite-containing vacuole and parasite destruction which we did not detect in infected SYNs at any timepoint tested. Our data show that SYNs are Toxoplasmacidal, and *T. gondii* that invade this specific trophoblast cell type are able to form what appear to be functional vacuoles, but are ultimately destroyed in a fashion reminiscent of some Toxoplasmacidal drugs (8, 28). Hallmarks of parasite killing are vacuolation of the parasites, breakdown of the vacuolar tubulovesicular network, and lack of integrity of the parasitophorous vacuolar membrane and leakage of host cytoplasmic contents into the lumen of the compromised vacuole. These data place PHT cells, and specifically the subpopulation of fused SYNs, into a rare class of cells that not only restrict the growth of *T. gondii* after invasion in the absence of any external stimuli, but actively destroy invaded parasites. While we did not compare them head to head with SYNs, neutrophils restrict *T. gondii* growth after invasion but are possibly less Toxoplasmacidal compared to SYNs given that neutrophils have been implicated in the spread of *T. gondii* throughout the intestine in a murine model (29). Head-to-head comparisons between SYNs and innate immune cell types like neutrophils will help to illuminate what potential killing mechanisms might be shared, or not, between these cell types.

In addition to resisting *T. gondii* infection, our data show that PHT cells robustly induce the chemokine CCL22 in response to infection by a Myr-1 dependent effector secretion mechanism. We do not know which cell types within the PHT preparation produce CCL22 after exposure to *T. gondii*, but given the dependence upon successful invasion on this response a good candidate cell type is the CYT rather than the SYN, although cell-specific analyses are required to address this directly. Importantly, the major inflammatory responses induced in PHT cells by *T. gondii* infection are not induced (or induced much more poorly) after infection with *N. caninum*, a near relative of *T. gondii* that does not successfully infect humans or rodents, suggesting that it may be a host and/or parasite adaptation that may impact disease outcome.

The precise role of CCL22 in human pregnancy is unknown, but maternal cells express CCL22 at low levels throughout pregnancy, with increased levels associated with miscarriage (12). Moreover, the induction of chemokines, including CCL22 and CCL17, are associated with preterm birth in humans (30) and in small animal models (31). The precise role played by CCL22 during *T. gondii* vertical transmission remains to be determined. However, exposure of PHT cells to recombinant CCL22 had no impact on parasite replication (**Supplemental Figure 3**), supporting the idea that it has no direct anti-parasitic activity. A likely scenario is that the induction of CCL22 is aimed at alerting the maternal immune system to placental infection, where it could play any number of roles in mediating the dialogue between maternal and fetal tissues, such as enhancing immune cell-mediated protection at the maternal-fetal interface, or in terminating the pregnancy should levels reach a specific threshold. Given that CCL22 levels are elevated in maternal serum during healthy, infection-free pregnancies (12, 13), CCL22 and its recruitment of regulatory T cells may also play a role in immune tolerance throughout gestation, which is modulated in response to infection.

Our findings provide important insights into the molecular and cellular pathways utilized by human SYNs at both late and mid stages of gestation to restrict *T. gondii* access to the fetal compartment. In addition, by characterizing the immunological pathways induced by *T. gondii* infection of SYNs, our findings have uncovered potentially novel biomarkers of infection severity that might have important roles in shaping the maternal systemic immune response. These findings provide an example of the signaling crosstalk that exists between the maternal and fetal compartments and the mechanisms by which this signaling is impacted by parasite-associated effectors. Taken together, these findings provide important insights into *T. gondii-*induced congenital disease that could lead to the design of novel therapeutics aimed at reducing congenital toxoplasmosis.

## MATERIALS AND METHODS

### Cell culture

All cell and tissue cultures were incubated at 37°C and 5% CO_2_ and all media were supplemented with 10% FBS and 50 µg/mL penicillin/streptomycin. JAR and HTR8 cells were grown in RPMI-1640 media (HyClone); BeWo cells in F-12K (Corning); and JEG-3 cells in Eagle’s Minimum Essential Medium (EMEM; Lonza). To induce fusion of BeWo cells, cells were treated with 10 µM forskolin for 24 hours, then washed with PBS before infection. Primary human trophoblast (PHT) cells were isolated from healthy, term-pregnancies, and were cultured as described previously(3, 9). PHT cells were cultured for ~48h prior to infection to allow for SYN formation. Primary placental fibroblasts were isolated and cultured as described previously(32).

### Midgestation Placental Explants

Human placental tissue from less than 24 weeks gestation was obtained from the University of Pittsburgh Health Sciences Tissue Bank through an honest broker system after approval from the University of Pittsburgh Institutional Review Board and in accordance with the University of Pittsburgh anatomical tissue procurement guidelines. Chorionic villi, fetal membrane, and decidua were dissected and cultured in DMEM/F12 (1:1) supplemented with 10% FBS, penicillin/streptomycin, and amphotericin B. For *T. gondii* infections, isolated tissue was infected immediately following isolation with 2.5x10^4^-1x10^7^ parasites for ~24hrs. For imaging, tissue was fixed in 4% paraformaldehyde and imaging performed as detailed below.

### Parasites

Type I (RH) and type III (CEP) *T. gondii* and *N. caninum* (NC-1) tachyzoites were used for this study. All parasites were maintained by continual passage in human foreskin fibroblast (HFF) cultures incubated at 37°C and 5% CO_2_ in DMEM supplemented with 10% FBS, 50 µg/mL penicillin/streptomycin, and 2mM glutamine. The YFP-RH was a gift from David Roos, and the RH-*MYR1*-KO and RH-*MYR1*-KO/complemented parasites were gifted by John Boothroyd. For infections, infected monolayers were scraped and syringe-lysed to release the tachyzoites. These parasites were then pelleted at 800 x *g* for 10 minutes, resuspended in fresh media, filtered through a 5 µm filter, and counted to determine the appropriate dilution for infection. Mock inoculum was produced by filtering out the tachyzoites with a 0.2 µm filter.

Parasite growth curves were generated by luciferase assay (Promega) using luciferase-expressing CEP parasites. Briefly, at each time-point, samples were lysed using the passive lysis buffer (Promega) and stored at −20°C until at least 8h past the last time-point collection. Samples were then thawed and incubated with substrate, and fluorescence was measured.

### RT-qPCR and RNAseq

RNA was isolated from cultures using the GenElute™ Mammalian Total RNA Miniprep Kit (Sigma) and the associated DNase digestion set (Sigma). Both a NanoDrop and an Agilent bioanalyzer were used to determine sample quality. Sequencing libraries were prepared from 0.2-0.9 µg of total RNA by the TruSeq Stranded mRNA Library Preparation Kit (Illumina). The Illumina NextSeq. 500 was used for sequencing. CLC Genomics Workbench 9 (Qiagen) was used to map the RNAseq FASTQ reads to the human reference genome (hg19). Differential expression analysis was performed using the Deseq2 package in R(33) using a significance cutoff of *P*adj < 0.01, unless specified otherwise. Analysis of mock and *T. gondii*-infected HFF cells was based on datasets previously published and deposited into the Sequence Read Archive (SRA): SRR2644999, SRR2645000, SRR2645001, SRR2645002, SRR2645003, and SRR2645004. Hierarchical clustering of log2 transformed RPKM data was performed using MeViewer TM4 software. Data were either clustered as is or linearly mean centered using Euclidian distance. Color scales were adjusted for presentation purposes. RNAseq data have been deposited in the NIH short read archive (accession numbers pending).

For RT-qPCR analyses, RNA was isolated as described above and cDNA generated using the iScript cDNA synthesis kit (Bio-Rad), followed by qPCR using a StepOnePlus Real-Time PCR System (ThermoFisher). The DC_T_ method was used to determine gene expression and normalized to the human actin C_T_ of each sample. Primer sequences were as follows: Actin**—** ACTGGGACGACATGGAGAAAAA (Forward, 5’-3’); GCCACACGCAGCTC (Reverse, 5’-3’). `CCL22`**—**GTGGTGTTGCTAACCTTC (Forward, 5’-3’); GGCTCAGCTTATTGAGAATC (Reverse, 5’-3’).

### Microscopy

Cell monolayers and placenta explants were fixed in 4% paraformaldehyde and permeabilized with 0.1% Triton X-100 in 1x PBS. Primary antibodies were incubated for 1h at room temperature, followed by washing, then secondary antibodies conjugated to Alexa Fluor (Invitrogen) fluorophores for 30min at room temperature. Following washing, cells/explants were mounted with DAPI-Vectashield (Vector Laboratories) and imaging performed on an Olympus FV1000 laser scanning confocal microscope, a Zeiss LSM 710, or an Olympus IX83 inverted microscope. In some cases, imaged were adjusted for brightness and contrast using Photoshop or Fiji/Image J. Image J was used for image analyses. Transmission electron microscopy was performed as described previously (9).

Reagents and antibodies used for immunostaining studies include Alexa Fluor 594 or 633 conjugated phalloidin (Invitrogen), cytokeratin-19 (Abcam), LAMP2 (Santa Cruz), SAG-1 (mouse monoclonal D61S; ThermoFisher).

## CCL22 ELISA

CCL22 ELISAs were performed with the human CCL22/MDC DuoSet ELISA (R&D Systems) as per the manufacturer’s instructions.

### Luminex

Conditioned media from cells was analyzed by multiplex luminex by the University of Pittsburgh Cancer Institute (UPCI) Cancer Biomarkers Facility: Luminex Core Laboratory that is supported in part by award P30CA047904.

### Statistics

All statistics were calculated using GraphPad Prism. Experiments were performed with independent preparations of PHT cells and second trimester villous explants. The data are presented as the mean ±SD. The individual statistical analyses and associated *P* values are described in the individual figure legends.

## Acknowledgements

We thank Judy Ziegler (Magee Women’s Research Institute) for technical support, the UPCI Luminex Core Facility for multiplex assays, and the UPCI Tissue and Research Pathology/Health Sciences Tissue Bank shared resource for human placental tissue, both of which are supported in part by P30CA047904. This project was supported by NIH R01-HD075665 [C.B.C. and Y.S.], NIH R01-AI114655 [J.P.B.] and T32-AI049820 [S.E.A.]. In addition, C.B.C. is supported by Burroughs Wellcome Investigators in the Pathogenesis of Infectious Disease Award.

---

**Supplemental Figure 1:**
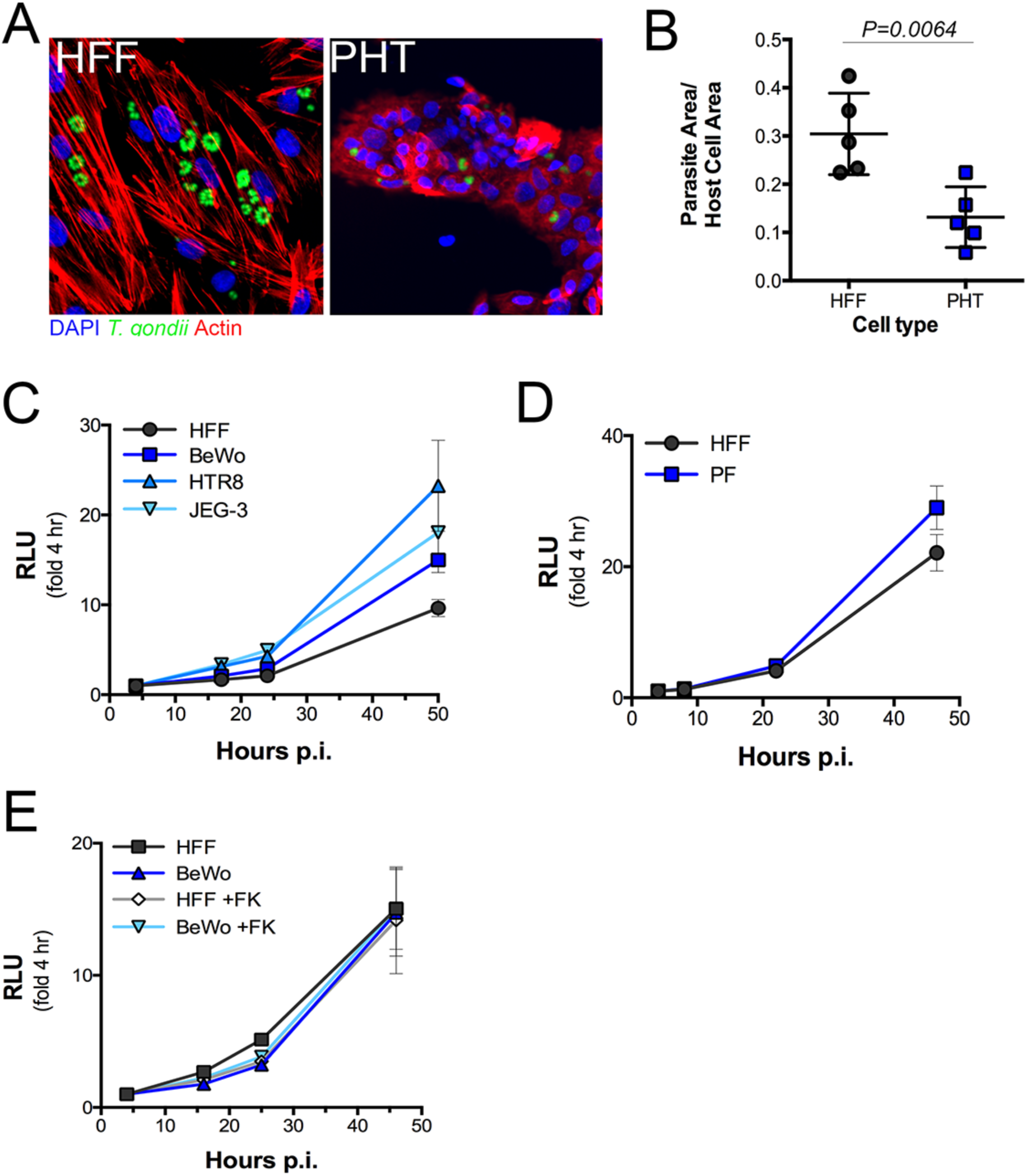
**(A, B)** Immunofluorescence microscopy of HFF and PHT cultures inoculated with T. gondii RH strain (green) at MOI= for 24h. (A) Representative images of HFF (left) and PHT (right) cultures. Actin is shown in red;; DAPI in blue. (B) Ratio of parasite to host cell areas based on immunofluorescence of five fields of view per culture. P=0.0064 based on 2-tailed T-ˇtest. (C, D) Growth curves of T. gondii CEP strain at MOI=0.5 in the indicated cell types, as measured by luciferase expression by parasites. Growth over time is indicated in relative light units (RLU) as normalized to expression at 4hpi;; and represented by the mean of three samples plus standard deviation. **(C)** T. gondii growth in three different trophoblast cell lines (BeWo, HTR8, and JEG-ˇ3) as compared to HFF cells. **(D)** Comparison of T. gondii growth in primary cultures of HFF and PF (placental fibroblasts). **(E),** T. gondii (CEP) growth in HFF and BeWo cultures +/-ˇ 10 μM forskolin pretreatment at MOI=0.5 as measured by luciferase expression by parasites. Growth over time is indicated in relative light units (RLU) as normalized to expression at 4hpi and represented by the mean of three samples plus standard deviation.

**Supplemental Figure 2.**
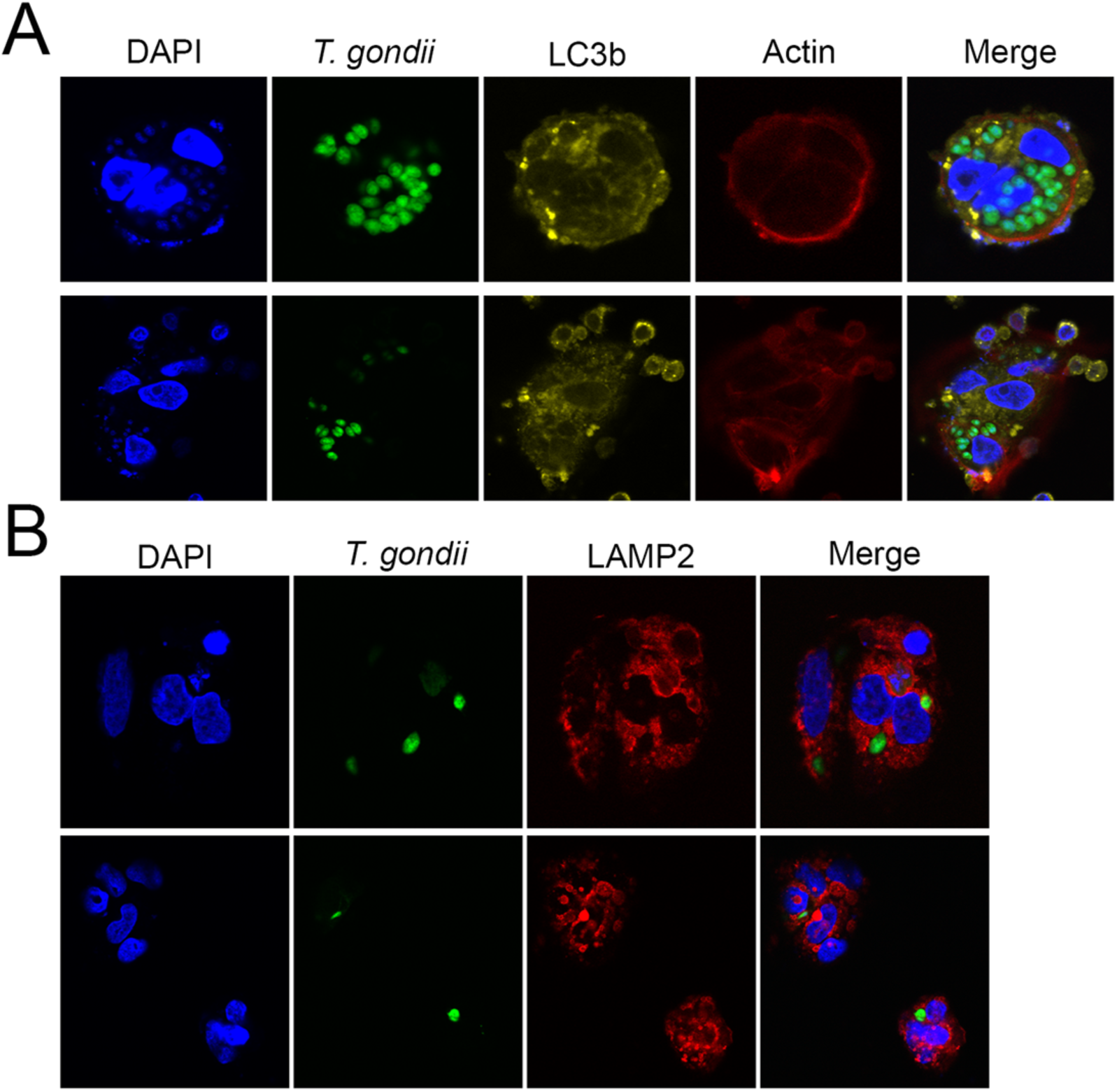
Immunofluorescence microscopy of PHT cells infected with *T. gondii* (YFP-RH, MOI=4) (green) In **(A),** LC3b staining is shown in yellow, actin in red, and DAPI-stained nuclei are shown in blue at 8rs p.i. In **(B),** lysosome-ˇassociated membrane protein 2 (LAMP2) is shown in red and DAPI is shown in blue at 24hrs p.i..

**Supplemental Figure 3.**
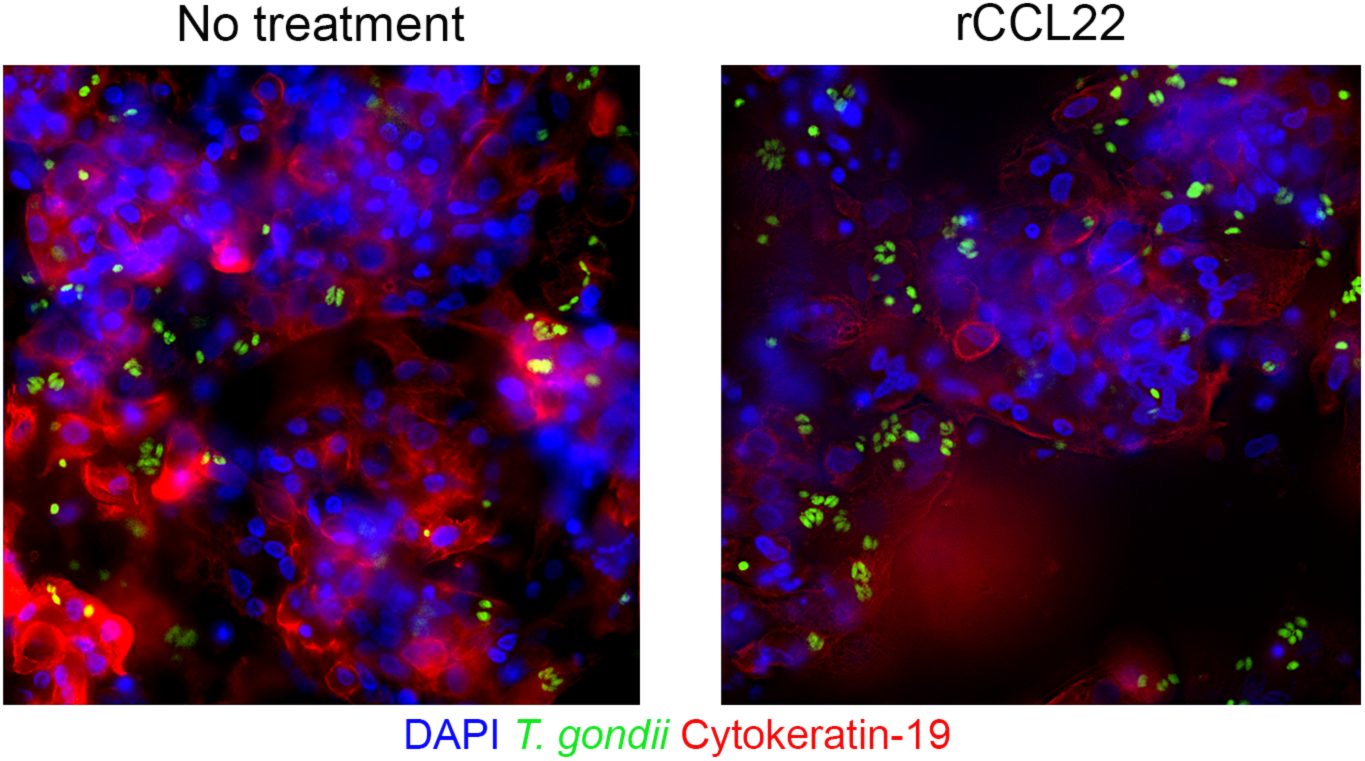
Immunofluorescence microscopy of PHT cells treated with 1ng/mL of recombinant CCL22 for 24hrs prior to infection with YFP-RH, MOI=3 for 24hrs. DAPI is shown in blue and cytokeratin-19 is in red.

## Literature cited

1. Torgerson PR & Mastroiacovo P (2013) The global burden of congenital toxoplasmosis: a systematic review. Bull World Health Organ 91(7):501–508.

2. Olariu TR, Remington JS, McLeod R, Alam A, & Montoya JG (2011) Severe congenital toxoplasmosis in the United States: clinical and serologic findings in untreated infants. Pediatr Infect Dis J 30(12):1056–1061.

3. Bayer A, et al. (2016) Human Placental Trophoblasts Produce Type III Interferons that Confer Protection Against Zika Virus Infection. Cell Host Microbe In press.

4. Delorme-Axford E, Sadovsky Y, & Coyne CB (2013) Lipid raft- and SRC family kinase-dependent entry of coxsackievirus B into human placental trophoblasts. Journal of virology 87(15):8569–8581.

5. Bayer A, et al. (2015) Human trophoblasts confer resistance to viruses implicated in perinatal infection. Am J Obstet Gynecol 212(1):71 e71–78.

6. Robbins JR, Zeldovich VB, Poukchanski A, Boothroyd JC, & Bakardjiev AI (2012) Tissue barriers of the human placenta to infection with Toxoplasma gondii. Infect Immun 80(1):418–428.

7. McConkey CA, et al. (2016) A three-dimensional culture system recapitulates placental syncytiotrophoblast development and microbial resistance. Science advances 2(3):e1501462.

8. Kamau E, et al. (2011) A novel benzodioxole-containing inhibitor of Toxoplasma gondii growth alters the parasite cell cycle. Antimicrob Agents Chemother 55(12):5438–5451.

9. Delorme-Axford E, et al. (2013) Human placental trophoblasts confer viral resistance to recipient cells. Proc Natl Acad Sci U S A 110(29):12048–12053.

10. Kafsack BF, Carruthers VB, & Pineda FJ (2007) Kinetic modeling of Toxoplasma gondii invasion. J Theor Biol 249(4):817–825.

11. Coffey MJ, et al. (2015) An aspartyl protease defines a novel pathway for export of Toxoplasma proteins into the host cell. eLife 4.

12. Freier CP, et al. (2015) Expression of CCL22 and Infiltration by Regulatory T Cells are Increased in the Decidua of Human Miscarriage Placentas. American journal of reproductive immunology 74(3):216–227.

13. Holtan SG, et al. (2015) Growth modeling of the maternal cytokine milieu throughout normal pregnancy: macrophage-derived chemokine decreases as inflammation/counterregulation increases. Journal of immunology research 2015:952571.

14. Franco M, et al. (2016) A Novel Secreted Protein, MYR1, Is Central to Toxoplasma's Manipulation of Host Cells. mBio 7(1):e02231–02215.

15. Braun L, et al. (2013) A Toxoplasma dense granule protein, GRA24, modulates the early immune response to infection by promoting a direct and sustained host p38 MAPK activation. The Journal of experimental medicine 210(10):2071–2086.

16. Reid AJ, et al. (2012) Comparative genomics of the apicomplexan parasites Toxoplasma gondii and Neospora caninum: Coccidia differing in host range and transmission strategy. PLoS Pathog 8(3):e1002567.

17. Landmann JK, Jillella D, O'Donoghue PJ, & McGowan MR (2002) Confirmation of the prevention of vertical transmission of Neospora caninum in cattle by the use of embryo transfer. Aust Vet J 80(8):502–503.

18. Davison HC, Otter A, & Trees AJ (1999) Estimation of vertical and horizontal transmission parameters of Neospora caninum infections in dairy cattle. Int J Parasitol 29(10):1683–1689.

19. English ED, Adomako-Ankomah Y, & Boyle JP (2015) Secreted effectors in Toxoplasma gondii and related species: determinants of host range and pathogenesis? Parasite immunology 37(3):127–140.

20. Bougdour A, et al. (2013) Host cell subversion by Toxoplasma GRA16, an exported dense granule protein that targets the host cell nucleus and alters gene expression. Cell host & microbe 13(4):489–500.

21. Shastri AJ, Marino ND, Franco M, Lodoen MB, & Boothroyd JC (2014) GRA25 is a novel virulence factor of Toxoplasma gondii and influences the host immune response. Infect Immun 82(6):2595–2605.

22. Hakansson S, Morisaki H, Heuser J, & Sibley LD (1999) Time-lapse video microscopy of gliding motility in Toxoplasma gondii reveals a novel, biphasic mechanism of cell locomotion. Mol Biol Cell 10(11):3539–3547.

23. Pfefferkorn ER (1984) Interferon-g blocks the growth of *Toxoplasma gondii* in human fibroblasts by inducing the host cell to degrade tryptophan. Proc. Natl. Acad. Sci. USA 81:908–912.

24. Pfefferkorn ER (1984) Interferon gamma blocks the growth of Toxoplasma gondii in human fibroblasts by inducing the host cells to degrade tryptophan. Proc Natl Acad Sci U S A 81(3):908–912.

25. Selleck EM, et al. (2013) Guanylate-binding protein 1 (Gbp1) contributes to cell-autonomous immunity against Toxoplasma gondii. PLoS Pathog 9(4):e1003320.

26. Choi J, et al. (2014) The parasitophorous vacuole membrane of Toxoplasma gondii is targeted for disruption by ubiquitin-like conjugation systems of autophagy. Immunity 40(6):924–935.

27. Foltz C, et al. (2017) TRIM21 is critical for survival of Toxoplasma gondii infection and localises to GBP-positive parasite vacuoles. Scientific reports 7(1):5209.

28. Lavine MD & Arrizabalaga G (2011) The antibiotic monensin causes cell cycle disruption of Toxoplasma gondii mediated through the DNA repair enzyme TgMSH-1. Antimicrob Agents Chemother 55(2):745–755.

29. Chtanova T, et al. (2008) Dynamics of neutrophil migration in lymph nodes during infection. Immunity 29(3):487–496.

30. Laudanski P, et al. (2014) Chemokines profiling of patients with preterm birth. Mediators Inflamm 2014:185758.

31. Arenas-Hernandez M, et al. (2016) An imbalance between innate and adaptive immune cells at the maternal-fetal interface occurs prior to endotoxin-induced preterm birth. Cell Mol Immunol 13(4):462–473.

32. Rausch K, et al. (2017) Screening Bioactives Reveals Nanchangmycin as a Broad Spectrum Antiviral Active against Zika Virus. Cell reports 18(3):804–815.

33. Love MI, Huber W, & Anders S (2014) Moderated estimation of fold change and dispersion for RNA-seq data with DESeq2. Genome Biol 15(12):550.

